# The MpsB protein contributes to both the toxicity and immune evasion capacity of *Staphylococcus aureus*

**DOI:** 10.1101/2021.06.30.450544

**Authors:** Edward J.A. Douglas, Seána Duggan, Tarcisio Brignoli, Ruth C. Massey

## Abstract

Understanding the role specific bacterial factors play in the development of severe disease in humans is critical if new approaches to tackle such infections are to be developed. In this study we focus on genes we have found to be associated with patient outcome following bacteraemia caused by the major human pathogen *Staphylococcus aureus*. By examining the contribution these genes make to the ability of the bacteria to survive exposure to the antibacterial factors found in serum, we identify three novel serum resistance associated genes, *mdeA, mpsB* and *yycH*. Detailed analysis of an MpsB mutant supports its previous association with the slow growing SCV phenotype of *S. aureus*, and we demonstrate that the effect this mutation has on membrane potential prevents the activation of the Agr quorum sensing system, and as a consequence the mutant bacteria do not produce cytolytic toxins. Given the importance of both toxin production and immune evasion to the ability of *S. aureus* to cause disease, we believe these findings explain the role of the *mpsB* gene as a mortality-associate locus during human disease.

## Introduction

*Staphylococcus aureus* is a major cause of human disease ranging from mild skin and soft tissue infections to more severe and life-threatening infections such as bacteraemia and pneumonia [1]. The continuing emergence of antimicrobial resistance is reducing therapeutic options and increasing the burden of this debilitating disease [2]. *S. aureus* bacteraemia (SAB) is associated with significant morbidity and mortality, where the 30-day mortality rate for SAB of approximately 20% has not substantially changed in over two decades [3]. This would indicate that improvements to infection control, surveillance and changes to therapeutic approaches over this time period have been insufficient in controlling this important health problem and suggests a greater understanding of this disease is needed. While SAB severity has been largely attributed to comorbidities of the host [4,5], our understanding of the role of specific bacterial factors in infection severity and disease progression is growing [6] and is the focus of this study.

To gain a deeper understanding of the role the bacteria play in the development and severity of SAB we previously performed a study of 300 SAB isolates, where we identified a number of genes as contributing to patient outcome that we refer to as mortality-associated loci (MALs) [6]. In this study we examine whether these MALs contribute to the ability of *S. aureus* to survive in human serum, given the importance of this activity to their ability to cause SAB. We identified three MALs as contributing to serum resistance, with one gene, *mpsB*, being of particular interest due to its association with the persister or small colony variant (SCV) phenotype associated with chronic *S. aureus* infections [7]. The MpsABC system participates in cation-translocation, and hence the generation of membrane potential, as well as CO_2_ transport [7,8]. Mutants of the operon, *mpsABC* exhibited a SCV-like phenotype and have been shown as attenuated in membrane potential and oxygen consumption rates [7]. MpsABC represents an important functional system of the respiratory chain of *S. aureus* that acts as an electrogenic unit responsible for the generation of membrane potential [7]. In this short communication we examine the contribution MpsB makes to serum resistance, demonstrate that it also contributes the ability of *S. aureus* to produce cytolytic toxins, where the effect the loss of this gene has on membrane potential is central to these key pathogenic capabilities.

## Methods

### Bacterial strains and growth conditions

A list of bacterial strains can be found in Table 1. All strains were cultured in Tryptic soy broth (TSB) for 18 h at 37^°^C with shaking. The transposon mutants were selected for using erythromycin (5 μg/mL). For the complemented pRMC2 strains, chloramphenicol (10 μg/mL) and anhydrous tetracycline (45-180 ng/mL) were added to the media [9]. CCCP (Cayman Chemical) was added to growth media at a final concentration of 0-6.25 µM where indicated.

**Table 1:** Strains used in this study.

| Strain | Description | Reference |
| --- | --- | --- |
| JE2 | USA300; CA -MRSA, type IV SCCmec; lacking plasmids p01 and p03; wild type strain of the NTML | [11] |
| JE2 RNAIII:*gfp* | JE2 transformed with a GFP tagged RNAIII fusion vector | [15] |
| *mpsB::tn* | *mpsB* transposon mutant in JE2 | [11] |
| *mpsB::tn* pRMC2 | *mpsB* transposon mutant in JE2 transformed with empty pRMC2 vector | This study |
| *mpsB::tn* p*mpsB* | *mpsB* transposon mutant complemented with *mpsB* gene cloned into the pRMC2 expression plasmid | This study |
| *mpsB::tn* RNAIII:*gfp* | *mpsB* transposon mutant transformed with a GFP tagged RNAIII fusion vector | This study |
| *agrA::tn* RNAIII:*gfp* | *agrA* transposon mutant transformed with a GFP tagged RNAIII fusion vector | This study |
| *hemB::tn* | *hemB* transposon mutant in JE2 | [11] |
| *crtM::tn* | *crtM* transposon mutant in JE2 | [11] |

### Genetic manipulation

The *mpsB* gene was amplified by PCR from JE2 genomic DNA using the primers *mpsB* FW 5’ atatagatctgaagaagtatttataggaggtgaaagg 3’ and *mpsB* RV 5’ tgaattcgagctcagatacttagcatcgcaacatatcatc 3’ and KAPA HiFi polymerase (Roche). The PCR product was cloned into the tetracycline inducible plasmid pRMC2 using *Bgl*II and *Sac*I restriction sites and T4 DNA ligase (NEB) creating p*mpsB*. This was transformed into RN4220 and eventually into JE2 *mpsB::tn* through electroporation.

### Serum resistance

Overnight cultures were standardised to an OD_600nm_ of 0.1 and incubated in 10% pooled human serum (Sigma Aldrich) diluted in PBS for 90 min at 37^°^C with shaking. Serial dilutions were plated on tryptic soy agar (TSA) to determine CFUs. The same number of bacterial cells inoculated into PBS, diluted, and plated acted as a control. Survival was determined as the percentage of CFU in serum relative to the PBS control. Relative survival was determined through normalisation to JE2.

### Minimum inhibitory concentrations (MICs)

MICs were performed according to the micro broth dilution method [10]. Briefly, overnight cultures were normalised to an OD_600nm_ of 0.1 in cation adjusted Mueller Hinton broth (MHB++) and 20 µL of resultant suspension used to inoculate 180 µL of fresh MHB++ containing gentamicin (Sigma). 1:2 dilutions were subsequently performed and incubated for 20 h at 37°C without shaking. The ability of the bacteria to survive the antibiotics was determined by quantifying bacterial growth (OD_600nm_) using a CLARIOstar plate reader (BMG Labtech).

### Antimicrobial peptide susceptibility

Antimicrobial peptide susceptibility Human neutrophil defensin-1 (hNP-1) (AnaSpec Incorporated, California, USA) and LL-37 (Sigma) susceptibility assays were performed as described previously [11]. Briefly, overnight cultures were normalised to an OD_600nm_ of 0.1 and incubated with 5 μg/mL of hNP-1 or LL-37 for 2 h at 37 °C. Serial dilutions were plated on tryptic soy agar (TSA) to determine CFUs. The same number of bacterial cells inoculated into PBS, diluted and plated acted as a control. Survival was determined as the percentage of CFU in serum relative to the PBS control. Relative survival was determined through normalisation to JE2.

### Carotenoid pigmentation

Carotenoid pigment quantification was performed as described previously with minor modifications [2]. Overnight bacterial cultures were used to inoculate 2 mL of fresh TSB in a 1:1,000 dilution which was subsequently grown for 24 h at 37°C with shaking (180 rpm). 850 μL of bacterial culture was centrifuged for 10,000 x g for 4 min, supernatant discarded and cells resuspended in 100% methanol. Cells were heated for 3 min at 55°C in a water bath, centrifuged at 10,000 x g for 2 min to remove cell debris and the extraction repeated twice. The absorbance of the methanol extracts was measured at 453nm using a CLARIOstar plate reader. JE2 *crtM::tn*, devoid of staphyloxanthin, was used as a negative control.

### Membrane potential

Overnight cultures were diluted to an OD_600nm_ of 0.05 in 20 mL of fresh TSB and grown to exponential phase. Cultures were subsequently normalised to 0.1 in 1 mL of PBS. Two sets of conditions were used for each strain. To one condition 10 µL of 500 µM CCCP was added. To both conditions 10 µL of 3 mM DiOC_2_(3) (ChemCruz) was added. Samples were incubated for 30 min at room temperature. Green fluorescence and red fluorescence were recorded using a CLARIOstar plate reader. Membrane potential was calculated according to the ratio of red: green fluorescence.

### Growth curve

Overnight bacterial cultures were diluted 1:1000 in fresh TSB. 200 µL of suspension was monitored for growth for 24 h at 37°C with 200 rpm shaking. OD_600nm_ readings were taken every 30 min using a CLARIOstar plate reader.

### Cytolytic activity

The monocytic THP-1 cell line (ATCC TIB-202) was used as previously described [13,14]. Briefly, cells were grown in 30 mL of RPMI-1640, supplemented with heat-inactivated fetal bovine serum (10 %), L-glutamine (1 μM), penicillin (200 units/mL) and streptomycin (0.1 μg/mL) (defined as complete medium) in a humidified incubator at 37°C with 5% CO2. For toxicity assays, cells were harvested by centrifugation at 400 x g and resuspended to a final density of 1-1.5 × 10^6^ cells/mL in tissue-grade phosphate buffered saline (PBS), typically yielding > 95% viability. These were then incubated for 10 mins in the harvested supernatant of bacteria grown for 18 h in TSB (at a dilution of 10% for the JE2 strains) and THP-1 cell death quantified by trypan blue exclusion.

### Agr activity

A plasmid containing the RNAIII promoter fused to the green fluorescent protein (GFP) was transformed by electroporation into JE2, *mpsB::tn* and *agrA::tn* [15]. Overnight cultures were diluted to an OD_600nm_ of 0.05 in fresh TSB. 200 µL of suspension was then monitored for GFP fluorescence (485_nm_ excitation / 520_nm_ emission/ 1000 gain) and OD_600nm_ over a period of 24 hours (readings every 30 min with 200 rpm shaking between readings) using a CLARIOstar plate reader.

### Statistics

Paired two-tailed student t-test or One-way ANOVA (GraphPad Prism v9.0) were used to analyse the observed differences between experimental results. A *p*-value <0.05 was considered statistically significant.

## Results

In previous work we identified a number of MALs, or genes associated with host mortality following SAB (Table 2) [6]. Given the importance of the ability to survive exposure to the many antibacterial factors found in blood to the ability of *S. aureus* to cause SAB, we hypothesised that some of these genes may contribute to this activity. To test this, we exposed an isogenic set of strains in which each of the 11 MALs were inactivated by transposon insertion to human serum (10%), and quantified their ability to survive relative to the wild type strain. Three MALs were found to significantly affect the ability of *S. aureus* to serum survival: *mdeA* which contributes positively to serum resistance, whereas *mpsB* and *yycH* contribute negatively to this (Fig. 1).

**Table 2:**
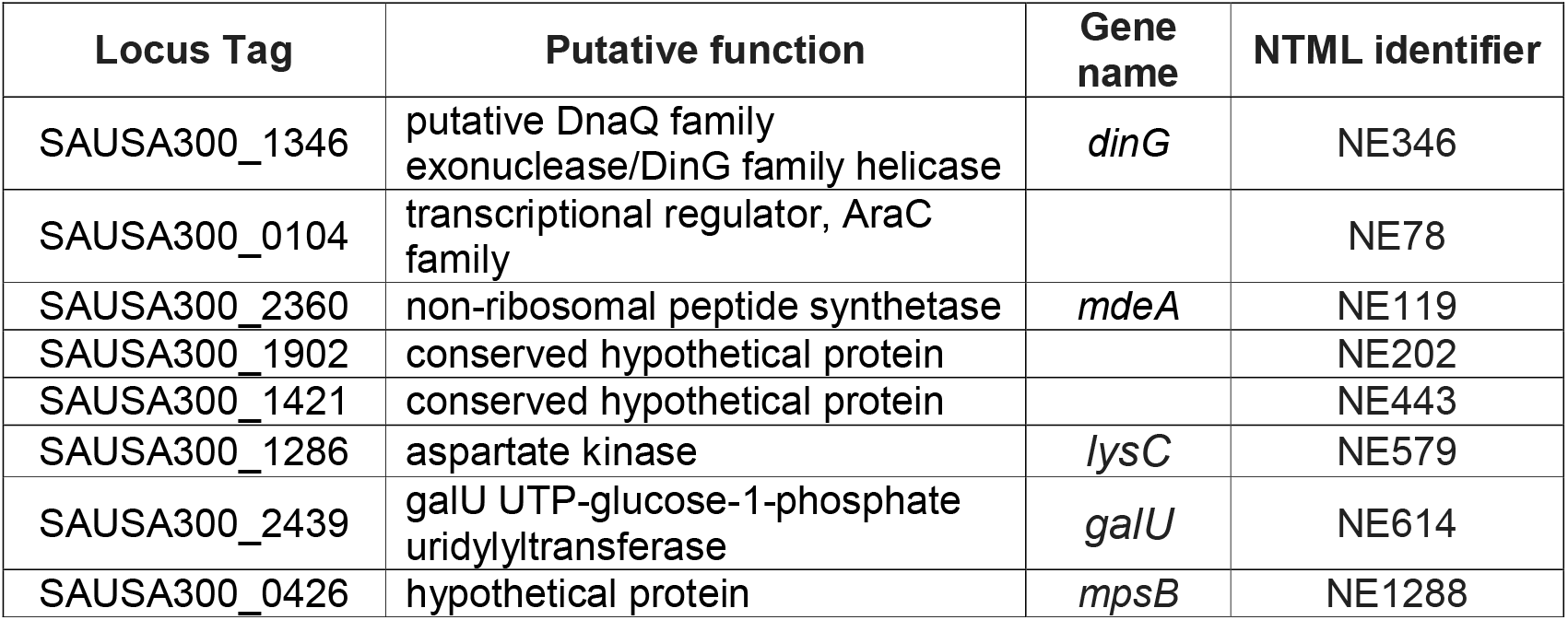
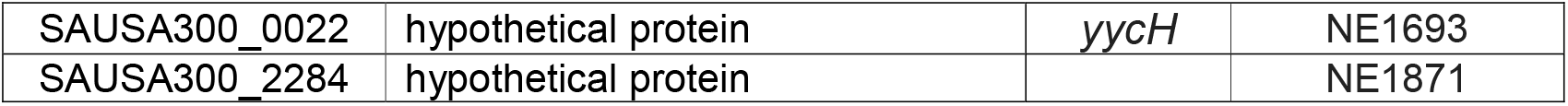
*S. aureus* genes previously associated with patient mortality (MALs).

| Locus Tag | Putative function | Gene name | NTML identifier |
| --- | --- | --- | --- |
| SAUSA300_1346 | putative DnaQ family exonuclease/DinG family helicase | *dinG* | NE346 |
| SAUSA300_0104 | transcriptional regulator, AraC family |  | NE78 |
| SAUSA300_2360 | non-ribosomal peptide synthetase | *mdeA* | NE119 |
| SAUSA300_1902 | conserved hypothetical protein |  | NE202 |
| SAUSA300_1421 | conserved hypothetical protein |  | NE443 |
| SAUSA300_1286 | aspartate kinase | *lysC* | NE579 |
| SAUSA300_2439 | galU UTP-glucose-1-phosphate uridylyltransferase | *galU* | NE614 |
| SAUSA300_0426 | hypothetical protein | *mpsB* | NE1288 |
| SAUSA300_0022 | hypothetical protein | *yycH* | NE1693 |
| SAUSA300_2284 | hypothetical protein |  | NE1871 |

**Figure 1:**
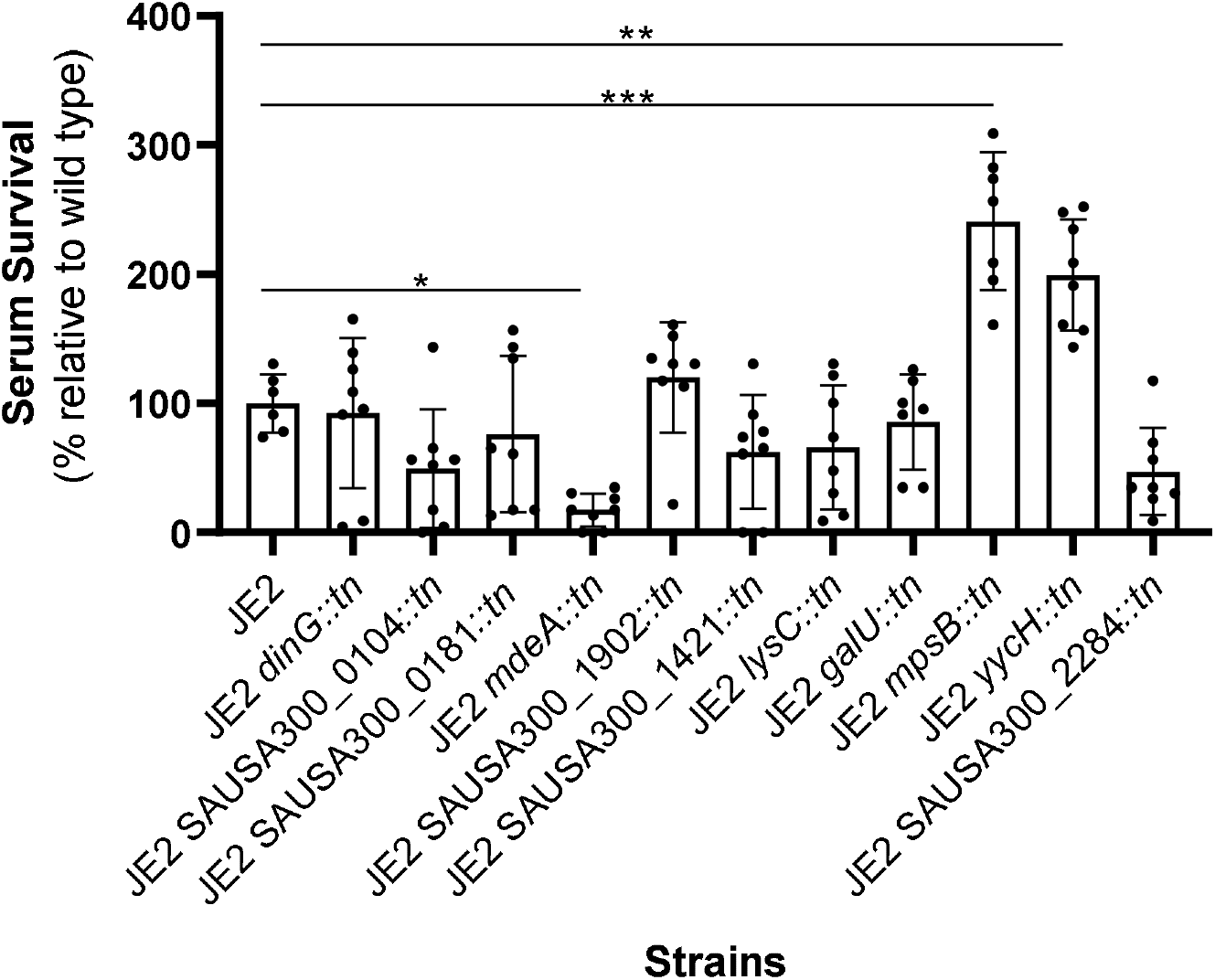
Serum survival screen of MALs. Mutants in which MALS were inactivated by transposon insertion were incubated in human serum for 90 minutes, serially diluted and plated on TSA for enumeration. Colony forming units (CFU) in serum were compared to CFU following a 90-minute incubation in PBS. The dots represent biological replicates, the bars represent the mean of the replicates, and the error bars the standard deviation. Statistics were performed using a one-way Anova. Significance was determined as <*0.01; **0.001; ***0.0001.

As part of the *mpsABC* locus, mutation of *mpsB* is likely to result in a change from the wild type to the persister or SCV phenotype [7]. SCVs are typically auxotrophic for menadione, hemin or thymidine, however, the Δ*mpsABC* mutant was found to be a CO_2_-dependant SCV and could be partially complemented using 5% CO_2_ [7,8]. To verify that our MpsB mutant displayed similar characteristics to those described previously we examined its growth on agar both with and without CO_2_ supplementation, where the colony size was smaller when grown in air compared to the wild type strain, but when grown in 5% CO_2_, the colony size of the mutant increased (Fig. 2a). We also examined whether carotenoid or staphyloxanthin biosyntheis was affected in the MpsB mutant, however, unlike other SCV types the MpsB mutant produced wild type levels of this protective pigment (Fig. 2b) [19]. We next examined whether membrane potential was affected in our MpsB mutant using the fluorescent stain DiOC_2_(3). This membrane potential indicator dye emits green fluorescence when it enters bacterial cells, but as it increases in concentration within cells that are maintaining membrane potential, the dye self-associates causing the fluorescence emission to shift to red [16]. Using this we found that the membrane potential of the MpsB mutant was significantly reduced to a level similar to that of a wild type strain exposed to carbonyl cyanide m-chlorophenylhydrazone (CCCP, a chemical that increases proton permeability thereby dissipating membrane potential) (Fig. 2c). [17]. The membrane potential of the MpsB mutant was also similar to that of a hemin auxotrophic SCV, where the *hemB* gene was inactivated (Fig. 2c). Another characteristic feature of SCVs is their enhanced resistance to the aminoglycoside antibiotic gentamicin [18]. To verify this for the MpsB mutant, we determined the minimum inhibitory concentration for gentamicin and the wild type JE2 strain, the MpsB mutant, the HemB mutant and the wild type strain when grown in CCCP, where the mutants and JE2 grown in CCCP were all more resistant to this antibiotic (Fig. 2d). Together these data suggest that the MpsB mutant displays some but not all the classic SCV phenotypic traits.

**Figure 2:**
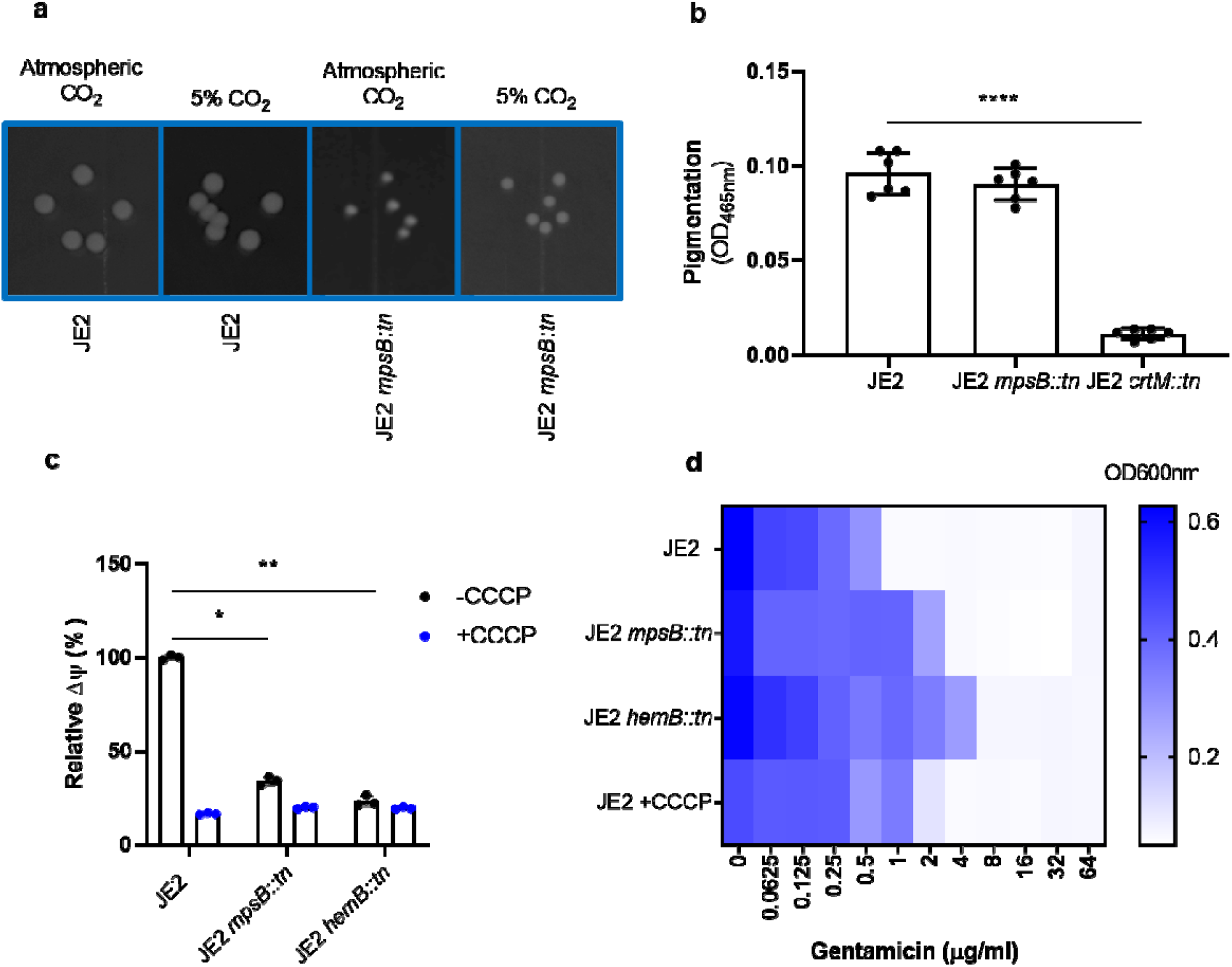
The MpsB mutant displays many SCV associated phenotypes. (**a**) Photograph of colonies of wild type *S. aureus* strain (JE2) and an isogenic MspB mutant grown with and without supplementation of 5% CO_2_, demonstrating the CO_2_ dependent effect on the growth of the mutant. (**b**) Staphyloxanthin biosynthesis is unaffected in the MpsB mutant when compared to the wild type stain. A staphyloxanthin biosynthesis mutant (JE2 *crtM::tn*) is provided as a negative control. (**c**) Membrane potential was measured using the fluorescent dye DiOC_3_ (3). The wild type JE2 and a *hemB* SCV (i.e. JE2 *hemB::tn*) are provided as controls where the native and post addition of CCCP membrane potential is shown. The MpsB mutant displays a similar reduction in membrane potential to the *hemB* SCV mutant. (**d**) Loss of membrane potential is associated with increased gentamicin resistance. When *mpsB* and *hemB* are inactivated the gentamicin MIC increases to 4 μg/ml and 8 μg/ml respectively as shown by an OD600nm heat map. Growth of JE2 in 0.5 μg/ml CCCP phenocopies the *mpsB:tn* and *hemB::tn* mutants and results in an increased gentamicin MIC. Significance was determined as <*0.01; **0.001; ****0.0001.

To ensure that MpsB alone was responsible for the observed increase in serum resistance we have observed for this mutant, and that there were no polar effects as a result of the transposon insertion into the *mpsB* gene, we cloned the gene and expressed it from an inducible promoter in the pRMC2 plasmid [9]. This complemented the serum resistance levels of the MpsB mutant back to wild type levels (Fig. 3a). Of the antibacterial components of serum, the antimicrobial peptides (AMPs), given their reliance on intact membrane potential for translocation, are most likely to be involved in the observed MpsB-associated sensitivity of the wild type strain [20–22]. To test this, we exposed the wild type and MpsB mutant to two AMPs HNP-1 and LL-37, and in both cases the MpsB mutant was more resistant to these molecules when compared to the wild type strains (Fig. 3b & c).

**Figure 3:**
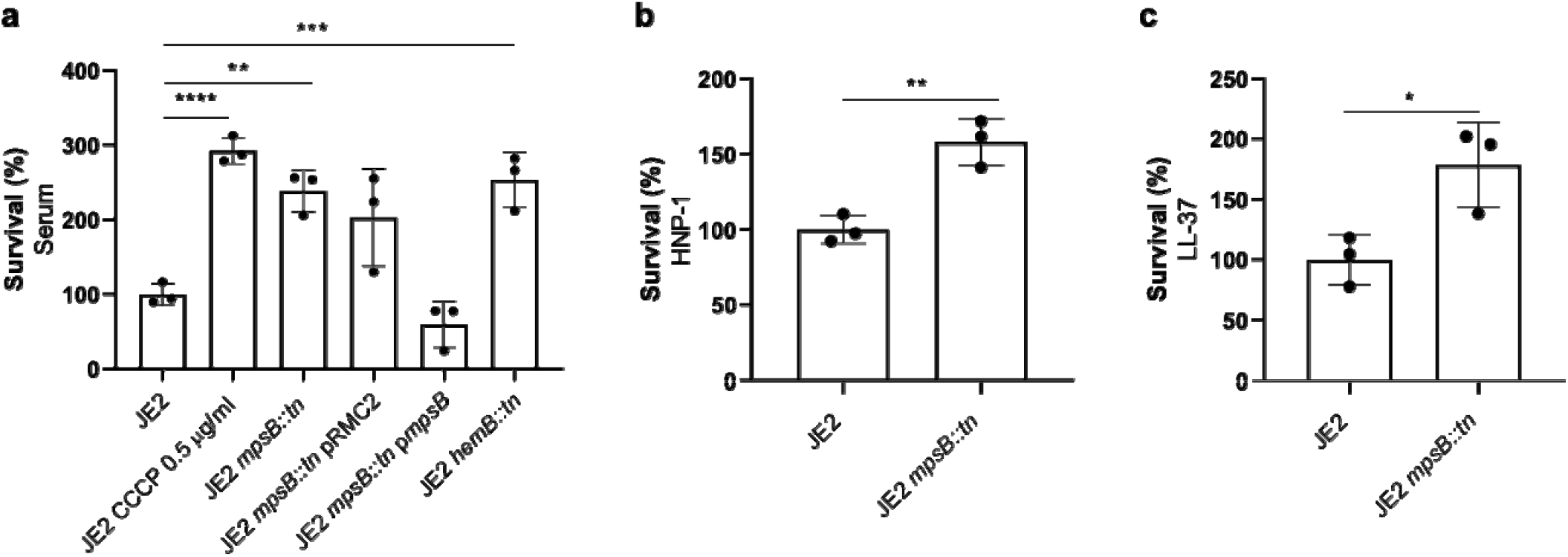
The resistance of the MpsB mutant to serum is mediated by their increased resistance to AMPs. (**a**) The effect the inactivation of *mpsB* has on serum resistance was confirmed by complementing the mutant with the *mpsB* gene expressed from an inducible plasmid (p*mpsB*). The serum resistance of the wild type JE2 exposed to CCCP and the *hemB* SCV are provided for comparison. (**b** & **c**) Loss of *mpsB* results in increased relative survival upon exposure to the AMPs HNP-1 and LL-37. The dots represent biological replicates, the bars represent the mean of the replicates, and the error bars the standard deviation. Statistics were performed using a one-way Anova and paired t-tests. Significance was determined as <*0.05; **0.01; ***0.001, ****0.0001.

In addition to serum resistance, mortality following SAB is significantly associated with the ability of the bacteria to secrete cytolytic toxins [6,23]. SCVs are frequently described as being non-haemolytic [24], and we examined here whether the MpsB mutant was impaired in its cytolytic capability. To test this, we incubated the bacterial supernatant with THP-1 cells, which are an immortalised monocyte progenitor cell line that are sensitive to the majority of cytolytic toxins produced by *S. aureus* [23]. We found the MpsB mutant to be less cytolytic, and this effect was complemented by expressing the gene *in trans* (Fig. 4A). The major regulator of toxin production for *S. aureus* is the Agr quorum sensing system [25], and to test how active this system is in the MpsB mutant we introduced a plasmid containing an Agr activity reporter fusion (RNAIII::GFP) [15]. We monitored both growth and RNAIII::GFP expression over a 24 h period and demonstrate the in a similar manner to and AgrA mutant, the Agr system does not get activated in the MpsB mutant, which explains the lack of cytolytic activity (Fig. 4B).

**Figure 4:**
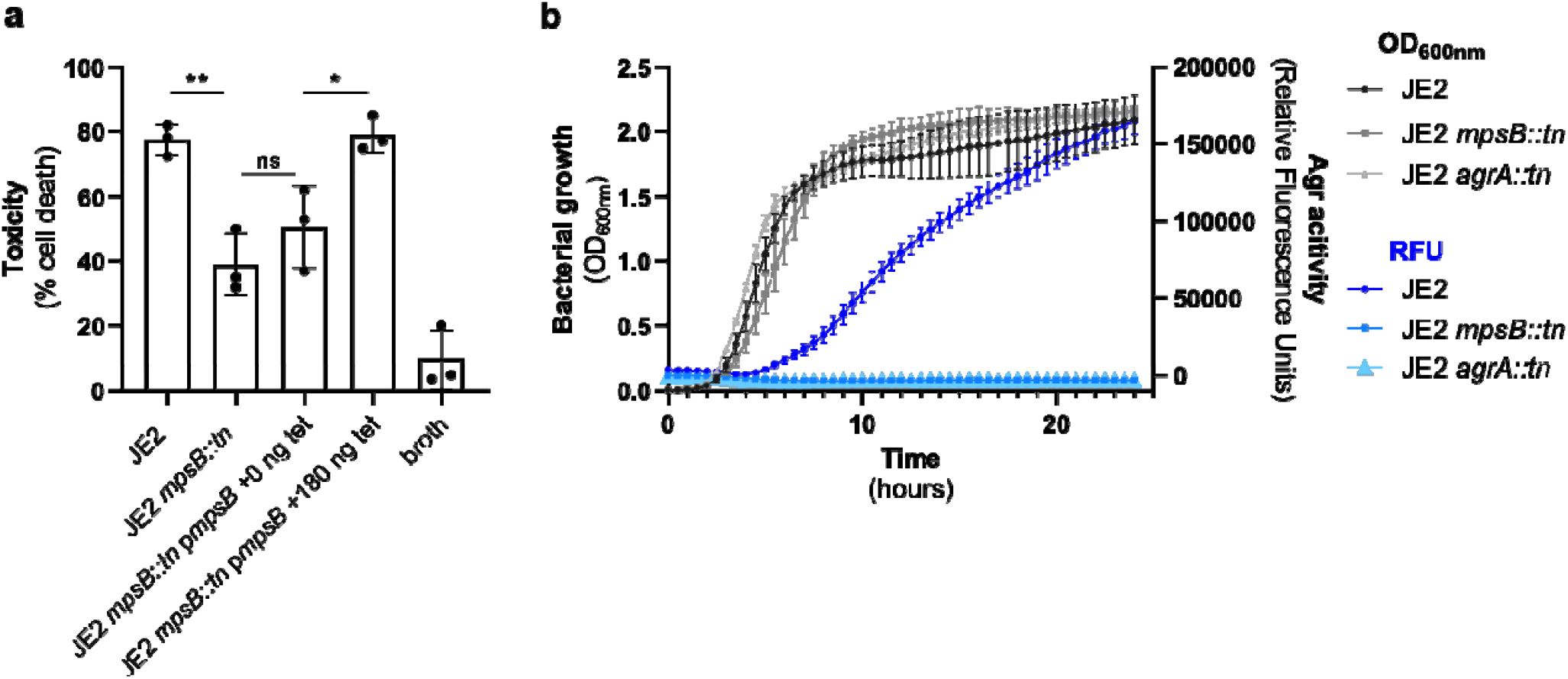
The MpsB mutant is less cytolytic mediated through a lack of activation of the Agr system. (**a**) Inactivation of *mpsB* results in a significant decrease in the ability of *S. aureus* to lyse the human monocyte (THP-1) cell line. This effect was complemented using a strain expressing a functional copy of *mpsB (*p*mspB)*. (**b**) Using an Agr fluorescence reporter system we found that when compared to the wild type strain JE2, the Agr system does not become activated in the MpsB mutant. An AgrA mutant is provided for comparison. The dots represent biological replicates, the bars represent the mean of the replicates, and the error bars the standard deviation. Statistics were performed using a one-way Anova and significance was determined as <*0.05; **0.01.

The activation of the Agr system is both density dependent, and highly sensitive to the metabolic status of the bacterium. The slow growth rate and effect of mutations in genes involved in key metabolic processes has been proposed as the reason why other SCVs don’t switch on their Agr system and remain non-cytolytic [19]. Interestingly, in this study while we see slower growth on agar (Fig. 1a) we don’t see any growth defects for the MpsB mutant when grown in broth (Fig. 4b), suggesting that a difference in relative density of the bacteria may not explain why the Agr system is not switched on. Given the critical role of two membrane proteins in the activation of the Agr system, AgrB and AgrC, we hypothesised a change in membrane potential could directly impact on their activity and cause the lack of activation we have observed for the MpsB mutant. To test this, we sought to identify a range of concentrations of CCCP that affect membrane potential, but doesn’t affect bacterial growth, which we found to be between 0.048 and 0.195 μM (Fig. 5a). We examined the effect these concentrations of CCCP have on Agr activation using a reporter system, where there were significant decreases in Agr activation (Fig. 5b). To ensure the effect of growth in CCCP on Agr activity was not due to effects on metabolic process throughout the growth curve, we also performed an experiment where we allowed the bacteria to grow to early stationary phase, monitoring Agr activation throughout. At hour 7, when the Agr system was becoming activated, we added the CCCP and found that this spike which would result in a sudden drop in membrane potential, had an immediate effect on Agr activity (Fig. 5c). We also performed THP-1 lysis assays on the bacteria following exposure to the low doses of CCCP, which confirmed that the effect we observed on Agr activity was sufficient to affect the cytolytic activity of the bacteria (Fig. 5d). This suggests that the Agr quorum sensing system requires a specific level of membrane potential for efficient activation.

**Figure 5:**
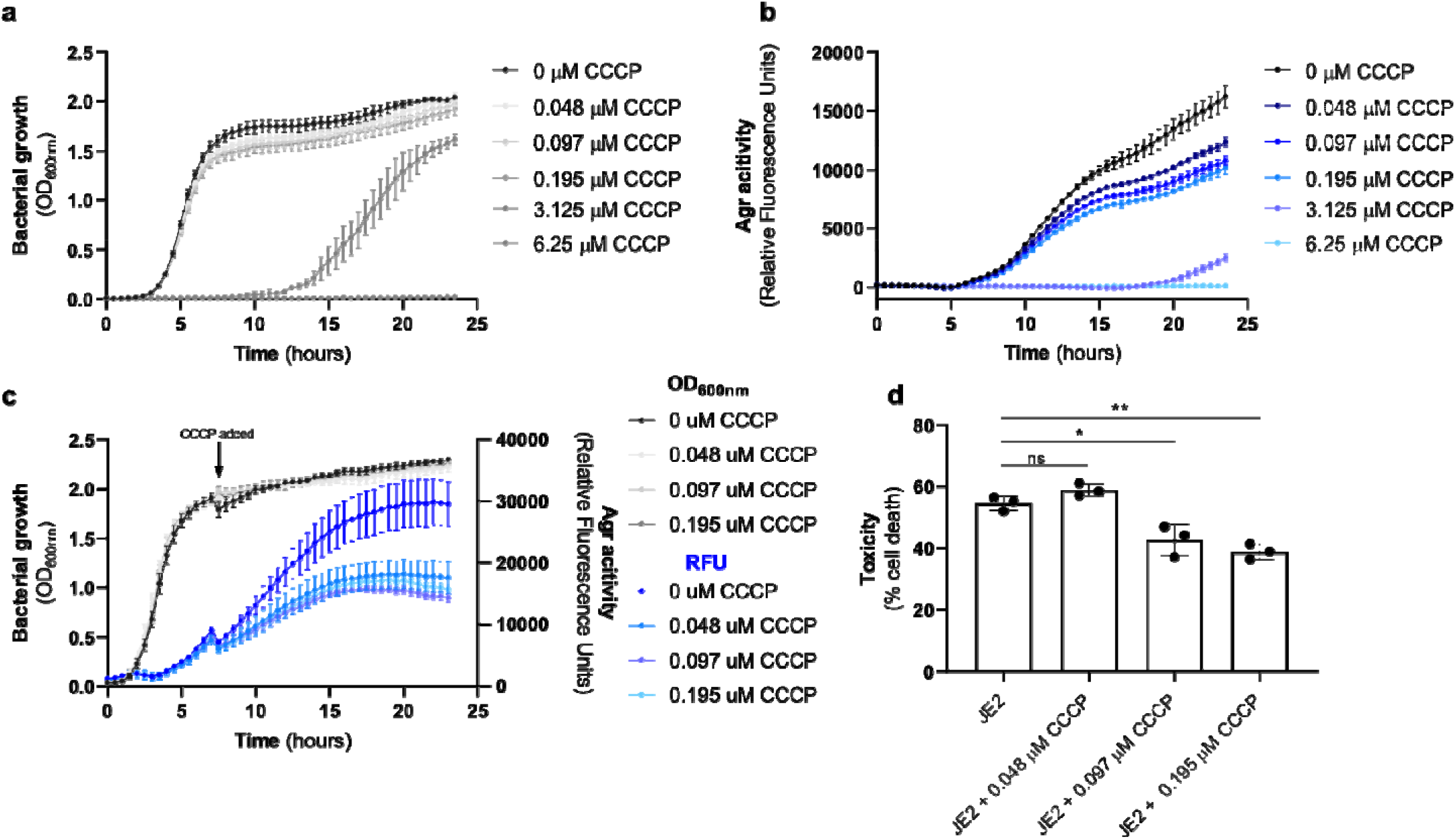
Activation of the Agr system require membrane potential. (**a & b**) The effect of a range of concentrations of CCCP on the growth (a) and Agr activity (b) of *S. aureus* was quantified over a 24h period, where concentrations ranging from 0.048-0.195 μM was found to affect Agr activation but not growth. (**c**) CCCP at a range of concentrations (0.048-0.195 μM) was spiked into cultures of bacteria as they reached stationary phase of growth, and the effect of this on Agr activity monitored. (**d**) The wild type S. aureus strain JE2 was grown in CCCP at a range of concentrations (0.048-0.195 μM) and the effect this has on the cytolytic activity of the bacteria quantified. The dots represent biological replicates, the bars represent the mean of the replicates, and the error bars the standard deviation. Statistics were performed using a one-way Anova and significance was determined as <*0.05; **0.01.

## Discussion

In this study we have identified three novel genes which contribute to the serum survival of *S. aureus*: *mdeA, yycH* and *mpsB*. We show that MdeA is involved in resisting the antimicrobial action of serum, whereas YycH and MpsB sensitize *S. aureus* to serum. MdeA is a chromosomally encoded efflux pump, that has been shown to confer resistance to a range of antibacterial compounds including quaternary ammonium compounds and the antibiotics novobiocin, mupirocin, fusidic acid and norfloxacin [26,27], and we hypothesise that MdeA may also expel antimicrobial agents found in serum. YycH is an auxiliary regulator of the WalRK two-component system, where a *yycH* mutant has been shown to have decreased WalRK activity leading to impaired autolytic activity [28]. It is currently unclear how loss of YycH causes increased serum survival, however, recent work has shown that alterations to cell wall architecture leads to increased survival in macrophages [29]. It is therefore possible that the antibacterial factors common to both macrophage and serum may be involved here.

Previous work on the Mps system has focused on either *mpsA* or the entire *mpsABC locus* and shown that the deletion of this locus results in an CO_2_ dependent SCV phenotype in the *S. aureus* HG001 background [7]. The SCV phenotype is widely accepted to be a strategy for protection against antibiotic therapy and aspects of the host immune system, and alongside their low virulence phenotype this makes them particularly well adapted for persistence. Here, we have focused on *mpsB*, and demonstrate that it directly contributes to the SCV associated phenotypes previously reported for *mpsA* and *mpsABC* mutants [7,8]. We show that disruption of *mpsB* in the JE2 background results in a partial SCV phenotype where we have a conditional effect on growth (slow on agar but normal in broth), reduced toxicity, increased resistance to the aminoglycoside gentamicin, to human serum and to antimicrobial peptides, but without the characteristic decrease in carotenoid pigment formation. Furthermore, we show that *mpsB* contributes to toxicity and the activity of the Agr system. A graphical summary of this has been provided (Fig. 6).

**Figure 6:**
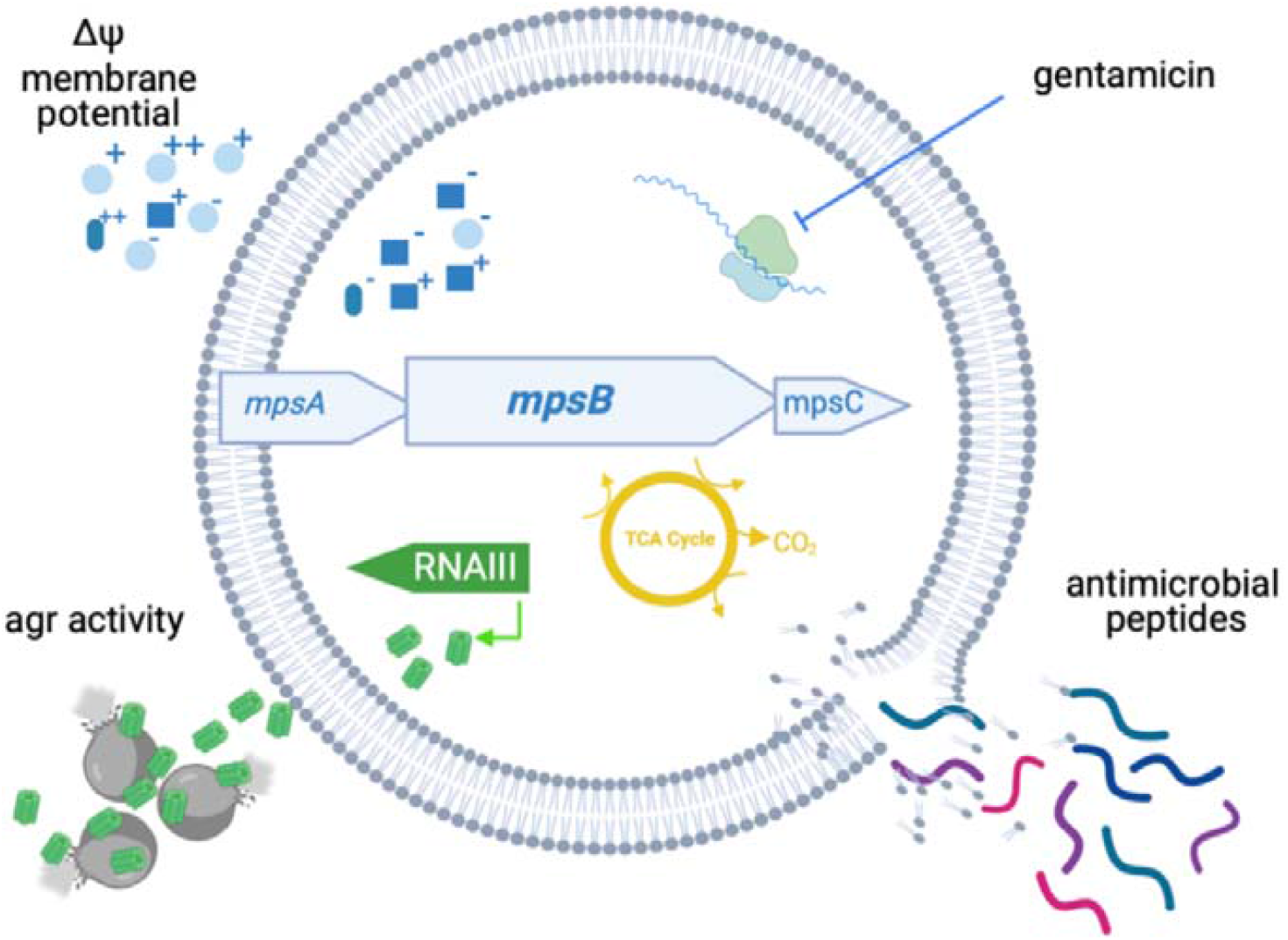
Graphical summary of the effects of the loss of MpsB. Disruption of *mpsB* disrupts membrane potential (upper left quadrant) which leads to increased resistance to the protein synthesis inhibiting antibiotic gentamicin (upper right quadrant) and cationic antimicrobial peptides (lower right quadrant). Membrane potential also effects the activity of the Agr system (RNAIII) leading to reduced cytotoxicity. Image created using *Biorender*.

The lack of toxin production by SCVs is well established, and is considered a result of a lack of activation of the Agr system due to the effects of the SCV conferring mutations on metabolism, and the associated reduction in cell growth. While the impact of *mpsB* inactivation was sufficient to switch off Agr activity, we did not observe an effect on growth in broth (Fig. 4b), which led us to consider whether membrane potential *per se* was directly involved in Agr activity. Using the membrane potential inhibitor CCCP, our findings suggest that this may be the case, as we see an immediate drop in Agr activity upon exposure to this chemical (Fig. 5c). While numerous factors have previously been shown to impact on Agr activity, a direct role for membrane potential has not been suggested prior to this study, where we hypothesise the effect may be mediated through the effect membrane potential may have on the activity of the two critical membrane located proteins of the Agr system, AgrB and AgrC.

By focussing on data from human infections, our aim is to unravel the pathogenic mechanisms utilised by *S. aureus* to cause disease. This approach has led to the identification of a number of MALs, and here we present the characterisation of the first three of these where their contribution to serum resistance is established. For *mpsB* we present further detailed characterisation of its activity, where its partial SCV phenotype allows it to survive the double edged-sword that human serum represent for bacteria - rich in metabolites while also highly immuno-protected [30,31]. While further work is underway to understand the role of the other MALs, this work provides an explanation for why the *mpsB* gene was associated with patient outcome following *S. aureus* bacteraemia.

## Authors Statements

### Contributions

EJAD developed the methodology, performed experiments, analyzed data and contributed to writing the manuscript. SD performed experiments, provided supervisory support, analyzed data, and contributed to writing the manuscript. TB developed the methodology, provided supervisory support, and contributed to writing the manuscript. RCM conceptualized the projects, provided supervisory support, analyzed data, and contributed to writing the manuscript.

